# A near-complete genome sequence of einkorn wheat provides insight into the evolution of wheat A subgenomes

**DOI:** 10.1101/2023.05.07.539731

**Authors:** Xiangfeng Wang, Hongna Li, Tao Shen, Xinrui Wang, Shu Yi, Tan Meng, Jie Sun, Xiaoliang Wang, Xiaojian Qu, Shisheng Chen, Li Guo

## Abstract

Einkorn wheat (*Triticum monococcum*) is one of the oldest cereal crops to be domesticated by human beings, playing essential role in early agriculture development. Today, it is considered an important genomic resource for modern wheat improvement, especially for resistance against pests and diseases. However, the exploration and utilization of useful genes from *T. monococcum* is limited due to the lack of a reference genome and annotation for this species. Here, we report a near-complete genome assembly for *T. monococcum* with a total length of 5.11 Gb with a contig N50 of 131.2Mb and scaffold N50 of 728.66Mb, representing a genome assembly of highest quality for any wheat genome reported. Phylogenomic analysis confirmed *T. monococcum* is closely-related to *T. urartu*, the progenitor of wheat A subgenomes. A 4AL/5AL terminal translocation is present in the diploid species *T. urartu* and *T. monococcum*, taking place before wheat polyploidization. *T. monococcum* has significantly expanded and unique gene families involved in DNA damage repair and heat stress tolerance, reflecting its adaptive evolution to cope with historical harsh climate in its natural habitat, South East Turkey. The genome sequence confirmed the introgression of *T. monococcum* rust resistance genes at 5A^m^L into modern bread wheat varieties. This near-complete reference genome of *T. monococcum* will be an essential resource for wheat functional and evolutionary genomic studies and expedite the cloning of useful genes in *T. monococcum* for future wheat improvement.

## INTRODUCTION

*Triticum monococcum* subsp. *monococcum* (2n = 2x = 14, A^m^A^m^), commonly known as einkorn wheat, is one of the oldest cereal crops to be domesticated around 12,000 years ago in South-East Turkey (Heun et al., 1997). This species played a significant role in the development of agriculture and was widely cultivated for centuries in the Middle East, Europe, Central Asia, and Africa before being replaced by free-threshing polyploid wheats. *Triticum monococcum* is closely related to *T. urartu* (genome A^u^A^u^), the donor of the A genome in durum and bread wheat (Dvorak et al., 1988). Today, cultivated einkorn is considered an important genomic resource for modern wheat improvement, especially for resistance genes against pests and diseases, including the stem rust resistance genes *Sr21* (Chen et al., 2018), *Sr22a*/*Sr22b* (Luo et al., 2022), *Sr35* (Saintenac et al., 2013), *SrTm4* (Briggs et al., 2015) and *Sr60* (Chen et al., 2020); the stripe rust resistance loci *QYrtm*.*pau-2A* and *QYrtb*.*pau-5A* (Chhuneja et al., 2008) and *Yr34*/*Yr48* (Chen et al., 2021); and the leaf rust resistance genes *LrTM16* (Sodkiewicz et al., 2008) and *Lr63* (Kolmer et al., 2010). However, the exploration and utilization of useful genes from *T. monococcum* is greatly limited due to the lack of a reference genome and annotation for this species. Here, we report a *de novo* assembly of *T. monococcum* genome by sequencing a cultivated accession PI 306540 collected in Romania and identified as having several stem rust resistance genes (Luo *et al*., 2022).

## RESULTS AND DISCUSSION

### Chromosome-scale genome assembly of *Triticum monococcum*

A total of 1189.92 Gb PacBio circular consensus sequencing (CCS or HiFi) reads, next-generation sequencing (NGS) reads and high-throughput chromatin conformation capture sequencing (Hi-C) reads by Illumina paired-end technology were generated for PI 306540, obtaining 36× (genome coverage) HiFi, 59× NGS and 138× Hi-C reads based on an estimated genome size of 5.09 Gb using k-mer frequency analysis of NGS read. Genome assembly of *T. monococcum* was first conducted from HiFi reads using *hifiasm* to yield a draft assembly of 5.11 Gb with contig N50 of 131.19 Mb. The HiFi assembly was then polished by *Pilon* using NGS reads and scaffolded to pseudochromosomes by *3d-DNA* and *Juicer* using Hi-C data. After genome polishing and decontamination of plastid and microbial sequences, the final assembly of *T. monococcum* is 5.11 Gb with a scaffold N50 of 728.66 Mb, containing seven chromosomes and 11 telomeres (Figure 1A). Mapping HiFi and NGS reads to *T. monococcum* assembly showed a mapping ratio of 99.87% and 99.57%, indicating a high completeness of assembly. Benchmarking Universal Single-Copy Orthologs (BUSCO) score of 96.4% using poales_odb10 database, LAI (LTR Assembly Index) score of 20.47 and Mercury QV score of 62.56 together suggest a high completeness and base-level accuracy for the *T. monococcum* genome assembly. Contiguity of this near-complete *T. monococcum* genome assembly (contig N50: 131.19 Mb) far exceeds those reported for *T. urartu* (contig N50: 0.344 Kb) and A subgenomes of tetraploid and hexaploid wheat genomes, representing the highest quality of wheat A subgenomes reported so far (Figure 1B).

**Figure 1.**
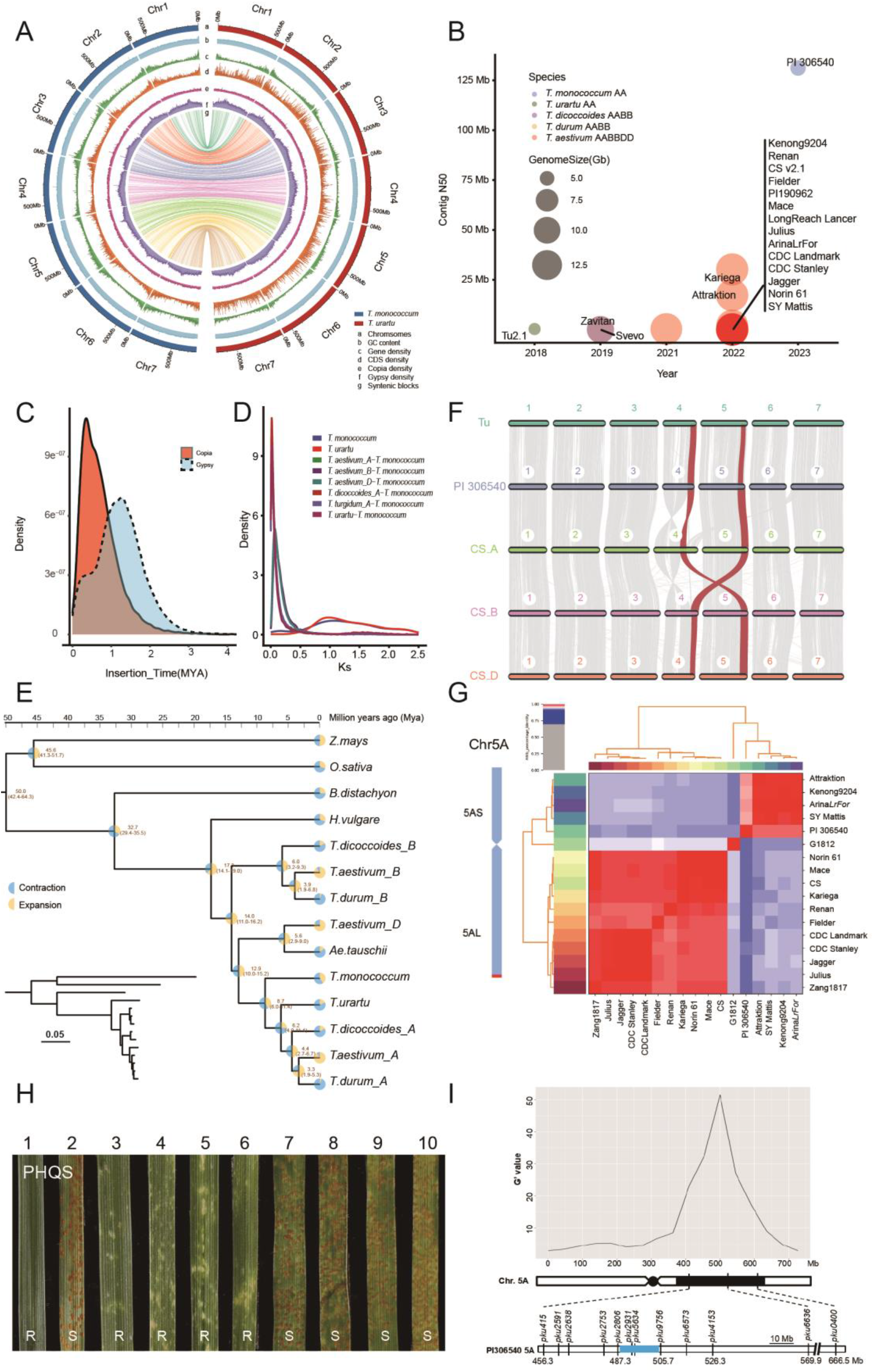
A near-complete assembly of *Triticum monococcum* genome illustrates the evolution of wheat A subgenomes. **(A)** Circos plot of genome features for *T. monococcum* (Tm) genome (this study) and published *T. urartu* (Tu) genome (Ling *et al*. 2018). Track a to g: chromosomes, GC content, gene density, CDS density, LTR/*Copia* density, LTR/*Gypsy* density and syntenic blocks of two genomes. **(B)** Comparison of genome assembly quality between *T. monococcum* and published wheat genomes. Each circle represents a genome. The fill colors represent different species. The diameter of the circle represents the genome size. **(C)** Synonymous substitution rate (Ks) density distributions of syntenic paralogs and orthologs. **(D)** Distribution of insertion times of *Gypsy* and *Copia* retrotransposons in *T. monococcum*. **(E)** A maximum likelihood phylogenetic tree of the genomes of major Triticeae species constructed using 2514 single-copy orthologs. **(F)** Syntenic blocks between the Tu, Tm (PI 306540) and the A, B and D subgenomes of bread wheat variety Chinese Spring (CS). Each line represents a syntenic block of five or more gene pairs with similarity of 80% or more. Red lines highlight representative chromosome rearrangements. **(G)** ANIb (average nucleotide identity blast) analysis of *T. monococcum* and other published reference genomes in the distal region of chromosome 5AL. This figure was generated using *pyani* with default settings and visualized using the pheatmap package in R. **(H)** Leaf rust reactions to *Puccinia triticina* race PHQS (isolate D3-2-182) inoculated on leaves of segregating resistant and susceptible plants from the cross of PI 119435 × G3116. 1, PI 119435 (*LrPI119435*); 2, G3116; 3-6, resistant F_3:4_ plants; 7-10, susceptible F_3:4_ plants. R,resistant; S, susceptible. **(I)** BSR-seq analysis mapped a leaf rust resistant locus on chromosome arm 5A^m^L in *T. monococcum* accession PI 119435. Top: The G’ value at each SNP was calculated within a window size of 100 Mb to identify genomic regions that showed obvious G′ peaks, which indicate the existence of QTLs for leaf rust. Middle: Schematic representation of the chromosome 5A. Black rectangle indicates the mapping region. Bottom: Physical map of *LrPI119435* in the reference genome *T. monococcum* accession PI 306540. The candidate region is highlighted in blue.

### Genome annotation

Genome annotation revealed *T. monococcum* genome has a repeat content of around 85% which mainly consists of *Gypsy* (46.06%) and *Copia* (21.85%) long terminal repeat retrotransposons (LTR). Time estimation using molecular clock suggests a burst of LTR insertion in *T. monococcum* genome at around 2.5 million years ago (Mya) (Figure 1C). A total of 58,909 protein-coding genes were predicted in *T. monococcum* using *MAKER* pipeline combining *ab initio* prediction, homolog proteins and transcriptomic data (NGS and PacBio Iso-seq) generated from five tissues (leaf, root, stem, seed, and spike) and different growth periods. Among them, 46,966 genes were functionally annotated by eggNOG-mapper, of which 73.92% are expressed in at least one tissue using a threshold of transcripts per million mapped reads (TPM) >= 1 (S4). Additionally, a total of 32,406 noncoding RNAs predicted including 2,617 tRNA, 6,825 rRNA and 4,150 small RNAs. The average gene density of *T. monococcum* is around 2.8 genes per Mb with a higher distribution of genes towards ends of chromosomes (Figure 1A).

### Phylogeny and genome evolution of *T. monococcum*

Synteny analysis suggested that *T. monococcum* has not experienced any recent whole genome duplications (WGDs) besides ones shared by all Triticeae (Figure 1D). Phylogenomic analysis of *T. monococcum* and nine representative monocot species based on 2,514 single-copy orthologs showed that *T. monococcum* is closely related to *T. urartu*, the progenitor of wheat A genome (Figure 1E). The divergence of *T. monococcum* from *T. urartu* was estimated to occur at around 8.7 Mya, before their divergence from the other A subgenomes of emmer wheat (*T. dicoccoides* and *T. durum*) and bread wheat (*T. aestivum*) at 6.2 Mya (Figure 1E). Comparative genomics analysis revealed a strong collinearity between *T. monococcum* and *T. urartu* (diploid, AA) and A subgenome of hexaploid (*T. aestivum*, AABBDD) wheat (Figure 1F). Sequence alignment shows that the 4AL/5AL terminal translocation is present in the diploid species *T. urartu* and *T. monococcum* and takes place before the polyploidization of wheat (Figure 1F).

*T. monococcum* has 1,753 expanded and 2,573 contracted gene families. The most significantly expanded gene family is ZF-GRF transcription factor (70 genes) involved in DNA damage repai. By contrast, bread wheat has fewer members of ZF-GRF gene family with many probably lost during domestication. In addition, *T. monococcum* has 997 unique gene families that are significantly enriched with functional terms such as heat tolerance conferred by genes encoding heat shock proteins, chlorophyll A-B binding proteins and protein kinases. *T. monococcum* is originated and mainly distributed in South East Turkey where crop plants frequently face harsh climate such as heat and drought stress conditions in history (Zaharieva and Monneveux, 2014). Therefore, the gene family expansion and unique gene family may reflect the adaptive evolution of *T. monococcum* to cope with drought and heat stress through mechanisms such as DNA damage repair.

### *T. monococcum* genome-assisted identification and characterization of rust resistance genes

Using the genome sequence obtained in this study, we confirmed the presence of the cloned stem rust resistance genes *Sr21, Sr22b*, and *Sr60* in *T. monococcum* PI 306540. In addition, the stripe rust resistance gene *Yr34* was located within a distal translocation of *T. monococcum* chromosome 5A^m^L into hexaploid wheat based on SNP analysis (Chen *et al*., 2021). To further explore the origin of the *Yr34* segment, the average nucleotide identity (ANI) metric was adopted to clarify genomic relationships between *T. monococcum* and other published reference genomes in the distal region of chromosome 5AL including *Yr34* (Figure 1G). We showed that four bread wheat varieties ‘Attraktion’, ‘Kenong9204’, ‘Arina*LrFor*’ and ‘SY Mattis’ may have the same distal 5AL *T. monococcum* translocation, which were clustered together with *T. monococcum* PI 306540 (Figure 1G). Within the *Yr34* candidate region in the PI 306540 genome, we detected one coiled-coil nucleotide-binding leucine-rich repeat (NLR) gene and six genes annotated as putative receptor-like protein kinases (RLKs) as potential candidate genes. In addition, we took advantage of the reference genome of PI 306540 obtained in this study and mapped a novel leaf rust resistance gene (temporary designation *LrPI119435*) in the *T. monococcum* accession PI 11943. The *LrPI119435* gene, conferring high resistance to Chinese *Pt* race PHQS (isolate D3-2-182, Figure 1H), was delimited within an 18.4Mb interval on chromosome arm 5A^m^L flanked by marker loci *pku2806* and *pku9756* (Figure 1I). The candidate region in PI 306540 includes several *NLRs* and *RLKs* that represent potential candidate genes. Among them, two genes (*TmPI3065405A01G042760* and *TmPI3065405A01G042780*) had significantly higher (*P* < 0.01) expression in the *Pt* (*Puccinia triticina*)-inoculated lines carrying the resistant allele compared to lines carrying the susceptible allele.

In summary, this newly constructed reference genome of *T. monococcum* represent the highest quality genome reference for wheat A subgenomes, which will serve as an essential genomic resource for functional and evolutionary genomic studies and expedite the cloning of useful genes in *T. monococcum* for future wheat improvement.

## METHODS AND MATERIALS

### DNA and RNA extraction

Einkorn wheat (*Triticum monococcum* subsp. *monococcum*) inbred line PI 306540 was grown in regular azalea pots filled with a combination of potting mix, clay and vermiculite in a greenhouse at Peking University Institute of Advanced Agricultural Sciences (Weifang, China). The fresh leaves of four weeks old einkorn wheat plants were harvested and washed with water and used for high molecular weight (HMW) DNA extraction using CTAB method (Porebski et al., 1997). Tissues of leaves, roots, stems, spikes, and grains of PI 306540 at different growing periods were collected and immediately frozen in liquid nitrogen and stored at -80 °C. Total RNA was isolated individually from collected *T. monococcum* tissues using Trizol (Thermal Fisher) agents following manufacturer recommendation protocol. The RNA Nano 6000 Assay Kit of Agilent Bioanalyzer 2100 system (Agilent Technologies, CA, USA) was used to evaluate the total RNA integrity.

### Genome sequencing

The high molecular weight DNA of *T. monococcum* was used as input to generate Illumina paired-end reads, PacBio HiFi reads and Hi-C (high-throughput chromatin conformation capture sequencing) reads. Paired-end short reads of *T. monococcum* were sequenced using Illumina NovaSeq 6000 to yield 301 Gb (∼59× genome coverage) data. To generate PacBio HiFi data, a total of 15 μg purified HMW genomic DNA were used to construct a standard PacBio SMRTbell library using PacBio SMRT Express Template Prep Kit 2.0 (Pacific Biosciences, CA). The CCS (circular consensus sequencing) was performed using a PacBio Sequel IIe instrument at Biomarker Technologies Corporation (QingDao, China) to generate 182Gb (∼36× genome coverage) HiFi reads with read length N50 of 18.2kb. Hi-C library construction was prepared from formaldehyde cross-linked chromatins of fresh young leaves using a standard Hi-C protocol (Belton et al., 2012). The constructed Hi-C sequencing library was first subjected to a test run of sequencing to evaluate valid interaction read pairs using *HiCPro* (v3.1.0) (Servant et al., 2015) before high coverage sequencing by Illumina NovaSeq 6000 to yield 705 Gb (∼138× genome coverage) paired-end reads.

### Genome assembly

The genome size and heterozygosity of *T. monococcum* were estimated using with *Jellyfish* (v2.3.0) (k-mer size = 21) (Marçais and Kingsford, 2011) and *Genomescope* (v2.0) (max k-mer coverage = 1,000,000) (Ranallo-Benavidez et al., 2020) from Illumina short reads, showing that the estimated genome size of *T. monococcum* is around 5.09 Gb. For genome assembly, we first assembled HiFi reads using *hifiasm* (v0.16.1) with the parameters of ”-l0” (Cheng et al., 2021). The resulting contigs were then aligned to the genome of chloroplast (GenBank: CM022232.1) and mitochondria (GenBank: MH051716.1) using *minimap2* (v2.24, -x asm5) (Li, 2018). Finally, we removed contigs from the assembly with at least 50% of their bases covered by chloroplast or mitochondria genome sequences (Naish et al., 2021). The assembled contigs were anchored using Hi-C data using the *Juicer* (v1.5) (Durand et al., 2016a) pipeline, with subsequent analysis by *3D-DNA* (v180419) (Dudchenko et al., 2017) and manual correction using *Juicebox* v1.11.08 (Durand et al., 2016b).

### Genome quality assessment

The obtained genome assembly was vigorously validated by three independent methods. Firstly, we performed LTR assembly index (LAI) assessment using *LTR_FINDER_parallel* (v1.2) (parameters: -harvest_out) (Ou and Jiang, 2019) and *LTR_retriever* (v2.9.0) (parameters: -inharvest) (Ou and Jiang, 2018). Secondly, the quality value (QV) was assessed using *Merqury* (v1.3) (parameters: k = 21 count) (Rhie et al., 2020). Additionally, the BUSCO (Benchmarking Universal Single-Copy Ortholog) was evaluated to reflect the completeness of genome assembly. The final *T. monococcum* genome assembly has a LAI of 20.47, QV of 62.56, and BUSCO score of 96.4%, suggesting the high accuracy and completeness of the assembly, respectively.

### Transcriptome sequencing and analysis

The total *T. monococcum* RNA was enriched the mRNA with polyA tail through Oligo (dT) magnetic beads. The mRNA was then subjected to transcriptome sequencing library construction using Illumina True-seq transcriptome kit (Illumina, CA) with an insert size of 370bp-420bp. The libraries were then sequenced by an Illumina Novaseq 6000 platform at Biomarker Technologies Corporation (QingDao, China) to generate 150bp paired-end reads. For full-length transcriptome sequencing (PacBio Iso-seq), about 5 μg mRNA was reverse-transcribed into full-length cDNA molecules with SMARTer™ PCR cDNA Synthesis Kit and the cDNA was further amplified by PCR. End repairing was conducted on amplified cDNAs, followed by SMRTbell adapter ligation. The ligation products are further treated by exonuclease to degrade failed ones before the Iso-seq library was sequenced using PacBio Sequal II instrument at Biomarker Technologies Corporation (QingDao, China). RNA-seq data was checked for quality using *fastp* software (v.0.23.2) (Chen et al., 2018), mapped to *T. monococcum* genome assembly using *hisat2* software (v.2.1.0) (Kim et al., 2019), followed by calculating mapping ratios by *samtools* (v.1.15). Iso-seq data was analyzed using CCS (v6.4.0) (https://ccs.how/; PacBio, 2021), *lima* (v2.6.0) (https://lima.how/; PacBio, 2021), *isoseq3* (v3.8.2) (https://github.com/PacificBiosciences/IsoSeq) and *pbmm2* (v1.9.0-2) (https://github.com/PacificBiosciences/pbmm2) and then used as input in genome annotation process.

### Genome annotation

For genome annotation, *ab initio* prediction and similarity alignment were firstly performed to annotate genome repeats via *RepeatModeler* (v1.0.11) (Flynn et al., 2020) and softmasked genome by *RepeatMasker* (v4.1.2.p1) (Nishimura, 2000). Gene model prediction combining *ab initio* prediction, homologous protein, RNA-seq and Iso-seq evidence was constructed using the pipeline described as follows. For the *ab initio* prediction, we firstly trained the *GeneMark-ET* model using *BRAKER2* (v.2.1.6) (Brůna et al., 2021) and then trained the *SNAP* (Johnson et al., 2008) *semi-HMM* model using *MAKER* (v3.01.03) (Cantarel et al., 2008). For homologous protein evidence, we downloaded the protein data of bread wheat from NCBI GenBank. For transcriptome evidence, RNA-seq reads were firstly aligned to our genome assembly through *hisat2* and performed reference-based assembly and *de novo* assembly of transcriptomes by *Trinity* (v2.8.4) (Grabherr et al., 2011). Iso-seq data was analyzed using CCS (v6.4.0), *lima* (v2.6.0), *isoseq3* (v3.8.2) and *pbmm2* (v1.9.0-2). These above evidence are integrated by *MAKER* to predict the final gene model. *Rfam/Infernal* (v.1.1.4) (Griffiths-Jones, 2005) and protein-coding genes with an AED < 0.5 were kept. Finally, *cmscan* (v1.1.4) are used to infer genome-wide non-coding RNAs.

### Synteny and phylogenomic analysis

Orthologues and orthogroups in 11 monocot genomes were inferred using *OrthoFinder* v2.5.4 (Emms et al., 2019) with default values setting and ‘-M msa’ activated. The longest predicted protein of each individual gene was used as the representative input for the *OrthoFinder* analysis. *TrimAl* v1.4.12 (Capella-Gutiérrez et al., 2009) was used to remove poorly aligned regions of protein multiple sequence alignments. *RAxML* v8.2.12 (Stamatakis, 2014) was used to build maximum likelihood phylogenetic trees using the GAMMAJTT model, with maize and rice as outgroup. Divergence time estimates from *TimeTree* (www.timetree.org), a public database, were used for selecting the range of lower and upper uniform calibration priors. Using the species phylogeny and the differentiation time of known species in TimeTree as correction. The *CodeML* and *MCMCTREE* programs in the *PAML* v4.9 (Yang, 1997) were run to analyze amino acid substitution models and estimate species divergence times. The *CAFE5* (Mendes et al., 2020) was applied to infer gene-gain and -loss rates in each genome. The orthogroups generated by *OrthoFinder* were regarded as distinct gene families and provided as inputs for *CAFE5* analysis. The identified genes were subjected to GO (Gene Ontology) enrichment and KEGG (Kyoto Encyclopedia of Genes and Genomes) enrichment analysis, with the p-value of significant enrichment was set as < 0.05. The syntenic analysis was performed by *JCVI* (v1.1.19) (Tang et al., 2008). We identified synteny blocks by performing an all-against-all LAST search and chaining the hits with a distance cutoff of 20 genes. Additionally, we required each synteny block to have at least 5 gene pairs. Ks values of syntenic block genes were calculated using *ParaAT* (v2.0) (Zhang et al., 2012).

### Mapping population and phenotyping

*T. monococcum* ssp. *monococcum* accession PI 119435 displayed high resistance (ITs = 0; to 1-) to Chinese *Puccinia triticinia* (Pt) race PHQS (isolate D3-2-182), whereas wild *T. monococcum* ssp. aegilopoides accession G3116 is highly susceptible (ITs = 3+) to the same race. We evaluated 113 F_2_ plants from the cross PI 119435 × G3116 and their corresponding F_2:3_ families (∼25 plants per family) with Pt race PHQS. We further confirmed the phenotypes of the critical recombinants by challenging 25 F_3:4_ plants homozygous for the recombination with the same race. After inoculation, plants were grown in growth chambers at 22°C (day) and 20°C (night) with a 16h light/8h darkness photoperiod. Infection types (ITs) were recorded at roughly 12 days after inoculation using a 0-4 scale (DL and Kolmer 1989).

### BSR sequencing of and QTL Mapping

Based on the phenotypic data, seedling of 15 homozygous resistant and 15 homozygous susceptible F_2:3_ families were grown in a growth chamber for ∼2 weeks, and pooled tissues were collected to construct the resistant bulk (R-bulk) and susceptible bulk (S-bulk) for RNA extraction. Total RNA of the parents and two bulks was isolated using the Direct-ZolTM RNA Kits (Zymo Research Co., Ltd., Irvine, CA, USA). RNA quality was determined using a NanoDrop OneC spectrophotometer (Thermo Fisher Scientific, Waltham, MA, USA) and RNA sequencing were conducted using Illumina’s HiSeq X sequencing system. Raw reads were cleaned using *fastp* v0.23.2 (Chen *et al*., 2018) to trim off adaptors and remove low-quality reads. The clean reads from the parents and two bulks were separately mapped to PI 306540 genome using *STAR* v2.7.10a (Dobin et al., 2013). *Freebayes* v1.3.6 (Garrison and Marth, 2012) was used for SNP calling. QTLs were detected using *QTLseqr* (Mansfeld and Grumet, 2018) with default parameters. The G’value of each SNP was calculated within a window 100Mb to identify genomic regions with G’peaks as possible QTLs.

### Development of molecular markers

Based on the BSR-seq results, single nucleotide polymorphisms (SNPs) between the parental lines PI 119435 and G3116 were selected to develop markers in QTL candidate regions. The primers were designed with *Primer3* online tools (https://primer3.ut.ee/) to amplify the intronic regions from conserved genes that carry the target polymorphisms. The PCR amplicons were sequenced using the Sanger method to confirm the presence of sequence polymorphisms between parents. The detected polymorphisms were subsequently used to develop Cleaved Amplified Polymorphic Sequence (CAPS) or Insertion-Deletion (InDel) markers (Konieczny and Ausubel, 1993; Bhattramakki et al., 2002).

### Gene expression analysis

BSR-seq data were employed to evaluate differentially expressed genes (DEGs) between resistant (PI 119435 and R-bulk) and susceptible lines (G3116 and S-bulk). Counts of cleans reads were calculated using *Kallisto* (Bray et al. 2016). DEGs within the candidate interval of chromosome 5A were identified using *DESeq2* (Love et al. 2014) following the parameters of p-value < 0.05 and |log2-foldchange| > 1.

### Stripe rust resistance gene Yr34/Yr48 and the published wheat reference genomes

Previously, the durable stripe rust resistance gene Yr34/Yr48 was reported to be located within a distal segment (∼9.5 Mb) of the cultivated *T. monococcum* chromosome 5AmL translocated to chromosome 5AL in modern polyploid wheat (Chen et al. 2021). In this study, the distal regions of chromosome 5AL from the released reference genomes of hexaploid wheat accessions Attraktion, Kenong9204, Chinese Spring, Keriega, Renan, Fielder, Zang1817, Arina *LrFor*, SY Mattis, Norin61, Mace, CDC Landmark, CDC Stanley, Jagger, Julius, and diploid wheat accession G1812 (Kale et al. 2022; Shi et al. 2022; Zhu et al. 2021; Athiyannan et al. 2022; Aury et al. 2022; Sato et al. 2021; Guo et al. 2020; Walkowiak et al. 2020; Ling et al. 2018) were used for comparative analysis.

## AUTHOR CONTRIBUTIONS

LG and SC conceived and supervised the study. HL and XQ collected the samples, conducted DNA extraction and generated sequencing data. XFW, HL, XRW, SY, TM and JS performed genome assembly, annotation, transcriptome and evolutionary analysis. HL, TS and XL performed the BSR-seq analysis and QTL mapping. XFW and HL prepared figures and tables. LG, SC, XFW and HL wrote and revised the manuscript. All authors read and approved the final manuscript.

## FUNDING

This project is supported by National Key Research and Development Program of China (2022YFD1201300), the Provincial Natural Science Foundation of Shandong (ZR2021MC056 and ZR2021ZD30), National Natural Science Foundation of China (31970317) and startup funding from Peking University Institute of Advanced Agricultural Sciences. SC and LG are also supported by Young Taishan Scholars Program of Shandong Province.

## ACKNOWLEDGMENTS

The authors would like to thank Bioinformatics Platform at Peking University Institute of Advanced Agricultural Sciences for providing the high-performance computing resources.

## Notes

### Competing Interest Statement

The authors have declared no competing interest.

